# Phages carry orphan antitoxin-like enzymes to neutralize the DarTG1 toxin-antitoxin defense system

**DOI:** 10.1101/2024.07.11.602962

**Authors:** Anna Johannesman, Nico A. Carlson, Michele LeRoux

## Abstract

The astounding number of anti-phage defenses encoded by bacteria is countered by an elaborate set of phage counter-defenses, though their evolutionary origins are often unknown. Here, we discover an orphan antitoxin counter-defense element in T4-like phages that can overcome the bacterial toxin-antitoxin phage defense system, DarTG1. The DarT1 toxin, an ADP-ribosyltransferase, modifies phage DNA to prevent replication while its cognate antitoxin, DarG1, is an ADP-ribosylglycohydrolase that reverses these modifications in uninfected bacteria. The orphan phage DarG1-like protein, which we term anti-DarT factor NADAR (AdfN), removes ADP-ribose modifications from phage DNA during infection thereby enabling replication in DarTG1-containing bacteria. AdfN, like DarG1, is in the NADAR superfamily of ADP-ribosylglycohydrolases found across domains of life. We find divergent NADAR proteins in unrelated phages that likewise exhibit anti-DarTG1 activity, underscoring the importance of ADP-ribosylation in bacterial-phage interactions, and revealing the function of a substantial subset of the NADAR superfamily.

## Introduction

In the past several years, there has been an explosion in the discovery of bacterial phage defense systems^1^. Not lagging far behind have been the discovery of counter-defense mechanisms by which phages block these bacterial immune mechanisms^2^. Most phage counter-defense mechanisms described to date are non-enzymatic in nature. Such direct, but non-enzymatic defenses consist of phage counter-defense elements that function by directly blocking the bacterial defense system via binding of the defense effector protein^3,4^, titrating an intermediate signaling molecule^5,6^, or structurally mimicking the defense target^7–9^. These proteins rarely have homologs outside of phage genomes, and are typically highly specific for the defense system that they inhibit. In contrast, relatively few enzymatic counter-defense proteins have been identified^2^. These have been detected in the context of signaling based immune systems, in which the phage infection triggers production of a small molecule alarmone that then activates a bacterial defense mechanism. Phage counter-defenses that either degrade these signals or produce competing decoy small molecules have been identified^10–12^. Outside of these mechanisms, there have been only two reports of phage counter-defense proteins that covalently modify a protein target via acetylation or ADP-ribosylation^13,14^. Whether enzymatic counter-defenses are less common, or just more challenging to characterize, is not yet clear.

One family of toxin-antitoxin (TA) systems, the DarTG systems, were recently shown to provide robust phage defense^15^. TA systems are two-gene operons that encode a toxin, which typically inhibits growth of the bacterial cell, and a cognate, neutralizing antitoxin that prevents toxin activity under homeostatic conditions^16^. DarT1 and DarT2 are DNA ADP-ribosyltransferase toxins that modify single-stranded DNA (ssDNA), targeting guanosine and thymidine, respectively^17–19^. ADP-ribosylated DNA cannot be replicated, and in some cases also cannot be transcribed, thereby preventing phage replication^15^. Two different types of DarTG systems have been identified: DarTG1 systems, in which the antitoxin, DarG1, is a protein of the NADAR (NAD and ADP-ribose) superfamily^17^, and DarTG2 systems, in which the antitoxin DarG2 contains a macrodomain, another type of ADP-ribosylglycohydrolase domain that has been extensively characterized as an ADP-ribose eraser in eukaryotic cells^18^. The antitoxins display specificity, with each antitoxin only able to remove ADPr from the base modified by its cognate toxin. In addition to its ADP-ribosylglycohydrolase macrodomain, DarG2 proteins have a second, DarT2-interacting domain which contributes to their ability to neutralize the toxin^20,21^. The NADAR domain of DarG1 contains an N-terminal extension of unknown function, but has not been shown to interact with its cognate toxin^17^. The two systems defend against different sets of diverse phages with only some overlap in the targets^15^.

To date, three phage counter-defense strategies have been discovered for phage evasion of DarTG-mediated defense^15,22^. The SECϕ18 phage, normally susceptible to DarTG2w defense, was shown to acquire resistance by accumulating mutations in its DNA polymerase that enable phage replication of ADP-ribosylated DNA. The other two known anti-DarTG counter-defenses, termed AdfA and AdfB, were identified in *E. coli* phage RB69 and *Vibrio cholera* phage ICP1, respectively via escape mutant analyses. Escape mutants in both cases had acquired a single nucleotide polymorphism that appeared to enable an existing counter-defense element to neutralize the DarT toxin^22^. Both AdfA and AdfB are small proteins with no homology to known proteins families that seem to block DarT activity through an interaction with the toxin. In both cases, the DarT-neutralizing variants of the genes were found to naturally exist within related phages, suggesting that the inactive variant may have evolved to neutralize another DarT homolog. We previously found that while the T4 phage encodes an active DarT1-blocking AdfA allele, a T4 Δ*adfA* phage remains resistant to DarTG1, even though this protein restores RB69 infectivity of DarTG1 cells when ectopically expressed^15^. These data suggest the presence of a second, unknown anti-DarTG1 factor in T4 phage.

Here, we investigate the molecular basis for the resistance of T4 and most of its relatives of the *Tevenvirinae* subfamily to the DarTG1 phage defense system. Using an expanded set of *Tevenvirinae* from the BASEL phage collection^23^, we find that neither the presence of AdfA in the genomes of these phages, nor its allele type (active or inactive), correlate with DarTG1 susceptibility. Instead, all *Tevenvirinae –* with the exception of RB69 – are resistant to DarTG1 defense. By co-infecting with a sensitive and resistant phage and characterizing the resulting chimeric viruses that have acquired DarTG1 resistance, we identified a second anti-DarTG1 counter defense element that is conserved within the T-even subfamily of phages. This element, which we have termed <u>a</u>nti-<u>D</u>arT factor <u>N</u>ADAR (AdfN), is both necessary and sufficient for phage counter-defense. AdfN is a member of the NADAR super family, which also includes the DarG1 antitoxin, and we show that like DarG1, AdfN has enzymatic activity that allows it to remove ADP-ribose from DNA. We further find that other phage NADARs have similar enzymatic activity. Phylogenetic analyses indicate that phages have likely independently co-opted NADAR domain proteins multiple times from different bacterial or archaeal sources, and that these NADAR proteins function as orphan antitoxins to reverse the activity of a TA-associated DarT1 toxin.

## Results

### Phage crosses reveal that gp30.3 is an anti-DarTG1 counter-defense element

Previous work identified AdfA as an anti-DarTG1 protein encoded in some T-even viruses^15^. However, deleting *adfA*, which encodes a protein we showed was sufficient to overcome DarTG1 defense when provided *in trans* from the bacterial cell, had no impact on the susceptibility of T4 to DarTG1-mediated defense^15^. To better understand the basis for phage resistance, we obtained a larger group of related T-even phages from a publicly available collection of *E. coli* phages that can infect the laboratory MG1655 strain, the BASEL phage collection^23^. To determine the sensitivity to DarTG1, we measured the titer of each phage on a DarTG1 containing strain and compared it to a strain bearing only an empty vector to calculate the efficiency of plaquing (EOP). We found that all *Tevenvirinae* phages assayed (Bas35-Bas47, as well as the classic T-even phages T2, T4, T6), are fully resistant to DarTG1 defense with the exception of RB69, which is strongly blocked by DarTG1 (Figure 1A). When we performed searches for *adfA* in this group of 17 *Tevenvirinae* phages, we found that only 13 of the phages encode *adfA* homologs (Figure 1A, circles). Of these *adfA* homologs, ten encode a histidine at position 164 of this protein (red circles), which we previously demonstrated enables DarT1 neutralization^15^, while the two closest RB69 relatives, Bas46 and Bas47, encode the non-functional RB69-like variants with an arginine at that position (Figure1A, blue circles; Figure S1). Thus, neither the presence of *adfA* nor the *adfA* type correlates with sensitivity to DarTG1 defense. Taken together, these data demonstrate that the *Tevenvirinae* phages must encode additional anti-DarTG1 counter-defense elements.

**Figure 1:**
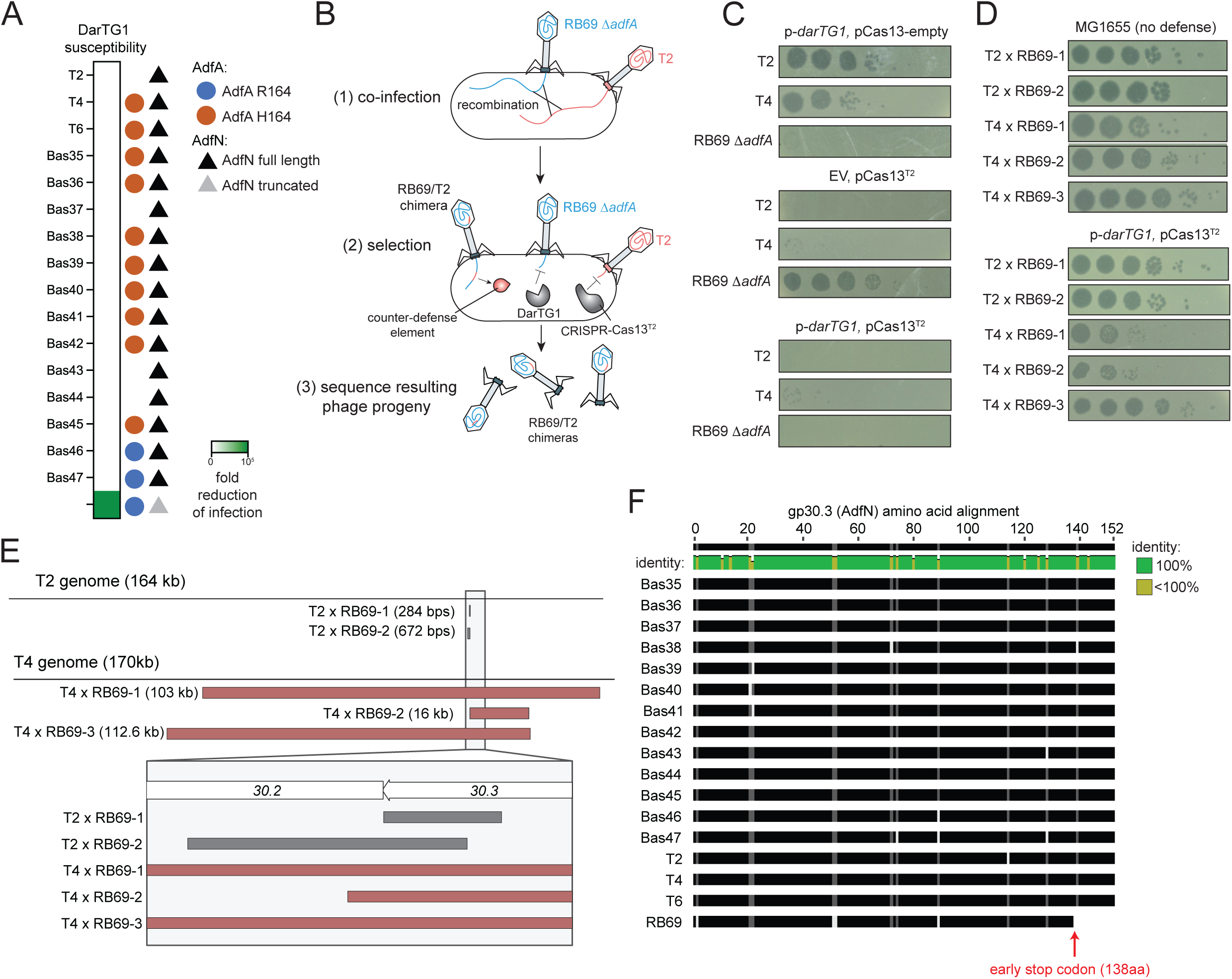
Recombination reveals the anti-DarTG1 counter-defense element in *Tevenvirinae*. (A) Efficiency of plaquing (EOP) for strains with DarTG1 compared to empty vector controls (left). The average of three independent replicates is presented. Circles indicate phage genomes that carry *adfA* genes. Blue circles encode the non-neutralizing variant of AdfA, while red circles indicate the functional, DarT1-neutralizing AdfA allele. Triangles indicate the presence of *adfN*. (B) Schematic of the co-infection and selection strategy for identifying the unknown counter-defense element. (C) Plaque assays with T2, T4 Δ*adfA*, RB69 Δ*adfA* on the indicated selection conditions with the indicated plasmid combinations. (D) Plaque assays of phage chimeras on a non-selective strain (upper) and double selection strain (lower). (E) Analysis of the T2 x RB69 Δ*adfA* and T4 Δ*adfA* x RB69 Δa*dfA* chimeras with the regions that transferred to RB69 depicted. The lower box is a zoomed region surrounding the genes *30.2* and *30.3* (F) Alignment of amino acid sequences of gp30.3 (AdfN) from the set of *Tevenvirinae* in our collection. See Figure S1 for a sequence level version of this alignment.

To identify additional counter-defenses, we took an approach that relies on the natural recombination between phage genomes that occurs when a bacterial cell is co-infected with two phages. We reasoned that in such a scenario, a subset of recombination events might result in the DarTG1-sensitive phage acquiring the DNA that encodes for a counter-defense element from the resistant phage (Figure 1B), similar to a previous study^24^. We selected RB69 as the DarTG1-sensitive phage, and T2 (which naturally lacks an *adfA* homolog) as the DarTG1-resistant, counter-defense-encoding host (Figure 1C). To avoid simply recovering AdfA^R164H^ mutants which arise very rapidly in RB69, we first generated RB69 Δ*adfA* for these experiments^15^. We co-infected a non-selective *E. coli* MG1655 strain with equal amounts of T2 and RB69 Δ*adfA* phages and collected the resulting lysate. To recover chimeric phages primarily derived from RB69, but bearing a T2-encoded counter-defense element, we programmed a CRISPR-Cas13 system to target T2, but not RB69 (pCas13^T2^)^25^ (Figure 1C). We then propagated the co-infection lysate on an *E. coli* strain expressing both the DarTG1 system and the CRISPR-Cas13 system, which blocks replication of both original phages (Figure 1C). Any plaques that arose on these plates were phages that were able to evade both DarTG1 and the T2-targeting CRISPR-Cas13. Resulting individual plaques were then selected, purified, and their resistance on the double selection strain was verified (Figure 1D). DNA from the chimeric phages was analyzed by Illumina sequencing (Figure 1E).

RB69 is sufficiently diverged from T2 such that distinctive patterns of single nucleotide polymorphisms (SNPs) can be used to distinguish the origin of DNA in the chimeric phages. We thus mapped the sequencing reads to both starting genomes, and using this method, were able to determine that the two RB69 x T2 chimeras analyzed were predominantly RB69, with only small regions transferred from T2 (Figure 1E). We performed the same experiment with a Δ*adfA* mutant of T4, a very close relative to T2, that we previously found is still resistant to DarTG1^15^ (Figure 1C-E). While the RB69 Δ*adfA* x T4 Δ*adfA* chimeras had transferred significantly larger regions than was the case in the RB69 Δ*adfA* X T2 chimeras, a comparison of five chimeric phages, two T2 x RB69 Δ*adfA* and three T4 Δ*adfA* x RB69 Δ*adfA*, revealed one core region that had transferred in each case, mapping to the 3’ region of the hypothetical gene *30.3* that is encoded within the entire set of T-even phages in our collection (Figure 1E). While *30.3* is also present in RB69, the *30.3* homolog in RB69 has a thymidine insertion at position 411 of the gene, which results in a frameshift and early stop codon in gene product 30.3 (gp30.3), truncating the protein 15 residues early (Figure 1F, S2A). Thus, the distribution of *30.3* among this phage group correlates with DarTG1 susceptibility: phages that encode full length versions are resistant to DarTG1, and the one phage with a truncated variant, RB69, is DarTG sensitive (Figures 1A, F). We renamed *30.3* to <u>a</u>nti-<u>D</u>arT factor <u>N</u>ADAR (*adfN*).

### AdfN (gp30.3) is necessary and sufficient to counter DarTG1 defense in T-even phages

To determine if AdfN is sufficient for resistance to DarTG1, we ectopically expressed the T4 AdfN in *E. coli* cells also expressing DarTG1 and measured phage defense against RB69. In the absence of AdfN, DarTG1 inhibits RB69 infection by about 5 orders of magnitude, but when AdfN^T4^ was provided ectopically, the ability of RB69 to form plaques on DarTG1-containing cells was fully restored (Figure 2A). We next tested whether this gene is responsible for the resistance of T-even phages to DarTG1. To this end, we made deletions of *adfN* from both T4 and T2 and found that in the absence of *adfN*, both phages become sensitive to DarTG1 defense, exhibiting a four-log reduction in plaques on lawns of DarTG1 compared to an empty vector (Figure 2B). We also made a T4 phage with both *adfA* and *adfN* deleted and found no additional change compared to the Δ*adfN* strain, suggesting that *adfN* is the primary anti-DarTG1 counter-defense factor in T4 (Figure S3A). We found that AdfN does not have the ability to counter DarTG2, whose toxin ADP-ribosylates the thymidine residue of DNA, as T4 Δ*adfN* is still DarTG2-resistant (Figure S3A), and overexpressing *adfN* ectopically does not rescue T5 from DarTG2-mediated defense (Figure S3B). Together, these data indicate that AdfN is a second anti-DarTG1 counter-defense element in the *Tevenvirinae*.

**Figure 2:**
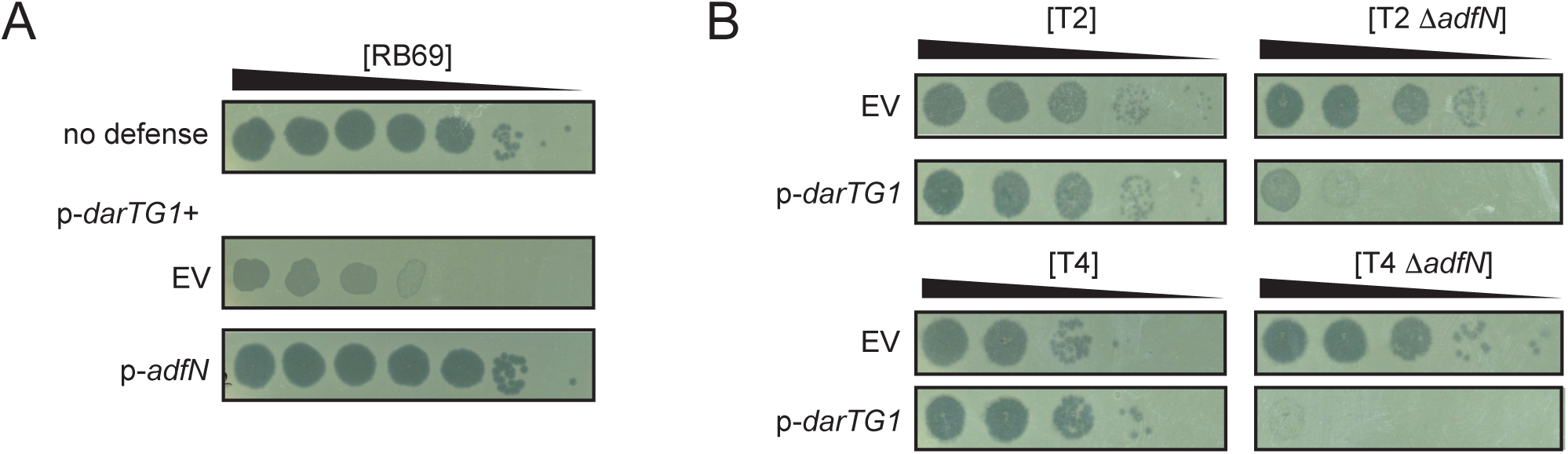
AdfN is necessary and sufficient for defense against DarTG1. (A) Plaques of RB69 on a strain without DarTG1 (upper) or with DarTG1 and either an empty vector (EV) or a vector encoding *adfN*. (B) Plaques of the parental or mutant phage on the indicated strains. All experiments were performed at least 3 times independently.

### AdfN is an DNA ADP-ribosylglycohydrolase

To gain more insight into how AdfN might be providing counter-defense, we next asked whether it associates with the DarT1 toxin in cells. In an earlier study, we found that AdfA^R164H^ neutralizes DarT1. Because AdfA has no apparent enzymatic domain or structural homology to any other proteins, we presumed that its mode of neutralization was likely to be via an interaction between the phage protein with the DarT1 toxin, just as type II antitoxins directly bind to and occlude the toxin active site^15,26^ though we had not formally tested whether these proteins associate. We hypothesized that AdfN might block DarT1 via direct interaction with the toxin, and thus directly assayed associations between DarT and these two counter-defense elements using a bacterial 2-hybrid assay. As wild-type *darT1* is toxic when expressed in *E. coli*, we fused a catalytically inactive variant of the toxin (with the mutation E152A, DarT*) to the T25 fragment of adenylate cyclase. Each candidate protein was fused to the T18 fragment of adenylate cyclase. Bacteria produce a blue pigment in the presence of 5-bromo-4-chloro-3-indolyl ϕ-D-galactopyranoside (X-gal) if the two proteins associate. Included as controls in this experiment were the non-functional, original AdfA from RB69 as well as its neutralizing, evolved variant identified previously (AdfA^R164H^)^15^. As expected, AdfA does not associate with DarT*, while AdfA^R164H^ displays a strong association, as indicated by the white and dark blue colors, respectively (Figure 3A). Consistent with the lack of a DarT1 binding domain in the DarG1 antitoxin protein, we also do not see evidence of an association between DarG1 and DarT1*, consistent with DarG1 acting as a type IV antitoxin, a class of antitoxin that does not directly interact with its cognate toxin. In contrast to AdfA^R164H^, we did not see evidence of a stable association between AdfN and DarT, suggesting that AdfN provides counter-defense through a different mechanism.

**Figure 3:**
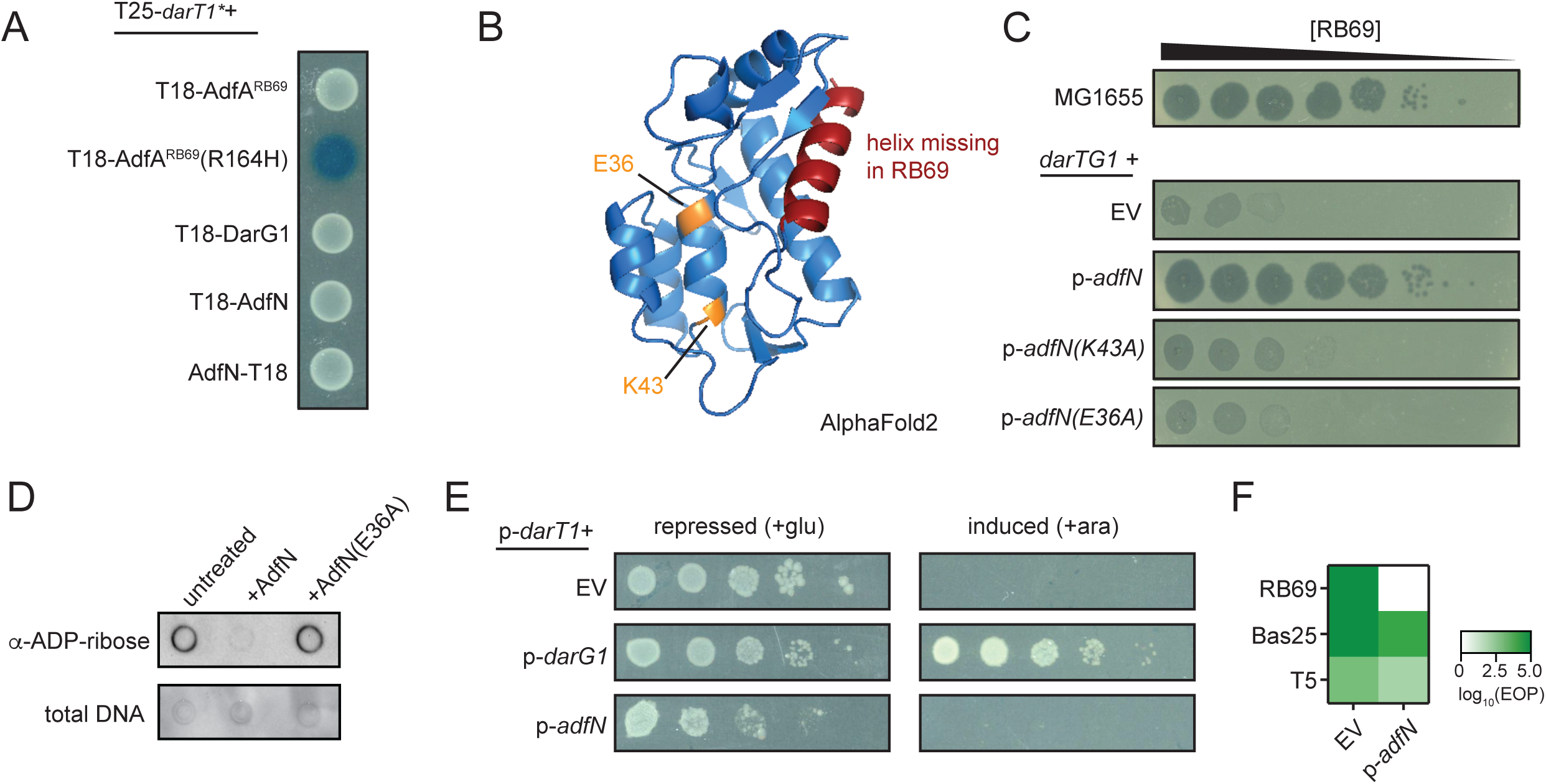
AdfN is a NADAR protein with DNA ADP-ribosylglycohydrolase activity. (A) Bacterial 2-hybrid experiment reporting on association of catalytically inactive DarT1 (DarT1*) bait with a series of prey proteins. Dark blue color indicates an association between the proteins; white means they do not associate. (B) AlphaFold2 model of AdfN, indicating a NADAR fold with conserved catalytic residues in orange, and the helix missing in the RB69 allele in red. (C) Plaques of RB69 in the presence of DarTG1 with ectopic expression of the indicated *adfN* variant. (D) ADP-ribosylation of DNA measured using a dot blot following an in vitro incubation with either AdfN or AdfN(E36A). (E) Colony forming units (cfu) of bacteria expressing DarT1 and either bearing an empty vector, a vector expressing *darG1*, or *adfN*. *darT1* was repressed with 0.2% glucose (left) or induced with 0.2% arabinose (right). (F) EOP data for the indicated phages on DarTG1/EV with either EV control or a vector encoding *adfN*. All experiments were performed independently at least 3 times.

To generate hypotheses regarding the function of the protein, we performed structural predictions of AdfN with AlphaFold2^27,28^ and searched for structural homology with the protein databank using DALI^29^ (Figure 3B). AdfN was predicted with high confidence (Z score > 9) to have structural homology to three structures of the NADAR superfamily, the DarG1 antitoxin from *Geobacter lovleyi*, a NADAR from *Phytophthora nicotianae* var. parasitica (NADAR^Pn^), and *E. coli* YbiA. NADAR proteins are DNA ADP-ribosylglycohydrolases that encompass the DarG1 antitoxin^17^, proteins involved in riboflavin metabolism, and many proteins of unknown function. We noted that the key conserved catalytic residues shown to be essential for NADAR activity in DarG1 and the *Phytophthora nicotianae* var. parasitica NADAR (*Pn*NADAR) are present in AdfN (Figure 3B, orange). We also noted that the early stop codon in RB69 would remove an entire terminal alpha-helix (Figure 3B, red). We thus hypothesized that, like DarG1, AdfN neutralizes DarT1 toxicity by enzymatically removing toxic ADP-ribose modifications from DNA. To experimentally test this hypothesis, we created two AdfN variants, each with a mutation in one of two key conserved catalytic residues, E36 and K43, that were shown in a structure of the PnNADAR to interact with ADP-ribose^17^. In contrast to AdfN, both the AdfN^E36A^ and AdfN^K43A^ were unable to rescue RB69 replication when expressed ectopically in the presence of DarTG1 (Figure 3C). We next purified a His-tagged variant of AdfN and incubated it with ADP-ribosylated DNA that had been purified from cells expressing DarT1, then measured ADP-ribosylation with an anti-ADP-ribose antibody in a dot blot. Confirming our hypothesis that AdfN is a DNA ADP-ribosylglycohydrolase, and consistent with another recent *in vitro* study of AdfN (gp30.3) activity^30^, we detected a strong signal from untreated DNA or DNA incubated with a catalytically inactive AdfN, but reduced ADP-ribose signal when the DNA had been incubated with AdfN (Figure 3D). Consistent with a recent report investigating AdfN *in vitro*^30^, these data demonstrate that, like the native DarG1 antitoxin, AdfN is an enzymatically active NADAR protein that detoxifies DarT1 through its ADP-ribosylglycohydrolase activity.

### AdfN is specific for activity in the context of *Tevenvirinae*

We next asked if AdfN, with its similarities to native DarG1 antitoxin, can stand in for DarG1. We first attempted to replace the native DarG1 antitoxin within a plasmid-encoded DarTG1 operon with AdfN, and were unable to clone this construct, suggesting that AdfN could not directly neutralize the toxin in *E. coli* under these expression conditions. We next asked whether ectopic expression of AdfN from a tetracycline promoter could restore the growth of cells expressing DarT1 from an arabinose promoter, which inhibits growth of *E. coli* in the absence of phage infection. Surprisingly, only the production of DarG1 – not AdfN – was able to restore growth of DarT1 expressing cells (Figure 3E). These data indicate that while AdfN is necessary and sufficient to counter DarT1 in the context of phage infection, surprisingly, it cannot function as an antitoxin in uninfected bacterial cells.

The knowledge that AdfN acts directly on DNA led us to hypothesize that even though AdfN can function on *E. coli* DNA *in vitro*, AdfN may have a strong preference for the DNA of *Tevenvirinae* phages *in vivo*. The DNA of these phages is hydroxymethylated and either glucosylated (Tequatroviruses, including T2, T4, and T6, to varying degrees)^31^ or arabinosylated (RB69)^32^. We hypothesized that AdfN may be adapted to interact with this heavily modified DNA, and may not be efficient at removing ADP-ribose from unmodified DNA, such as that found in *E. coli*. We predicted that if modified DNA is indeed important for AdfN activity, ectopic expression would not restore replication to phages with unmodified DNA in the context of DarTG1 defense. T5, in the *Markadamsvirinae* family, and Bas25, of *Queuovirinae*, are two phages inhibited by DarTG1 with no known DNA modifications. Consistent with AdfN preferring the DNA of *Tevenvirinae*, AdfN expression in *E. coli* provided only very modest rescue of less than a half a log of plaquing to T5, and no rescue to Bas25 (Figure 3F). We took a second approach to further decipher the requirement for AdfN activity. While it is not possible to generate a T4 phage with fully unmodified DNA due to the suite of T4 nucleases that degrade unmodified DNA, a T4 mutant lacking glucosylation (T4 Δ*agt* Δ*bgt*) has been generated. If AdfN requires glucosylation, we would expect that this phage would be unable to replicate in the presence of DarTG1. However, T4 Δ*agt Δbgt* exhibits normal replication on DarTG1 containing cells, indicating that AdfN has full activity on DNA that is only hydroxymethylated, and not further modified (Figure S3A, lower row). These results indicate that AdfN is most active in the context of the *Tevenviruses*, and that this specificity is likely due to hydroxymethylated DNA^15^.

### AdfN factors are found in diverse viral clades and have distinct substrate specificity

It has previously been shown bioinformatically that numerous phages encode NADAR proteins; indeed, the T4 gene encoding AdfN (*30.3*) was identified in this study, and a subsequent study demonstrated its ADP-ribosylglycohydrolase activity *in vitro*^17,30^. These phage NADARs cluster together, and we noticed that they are primarily found in phages that are part of the T4 superfamily^33,34^. While this group of NADARs form a cluster distinct from the DarG1-like (DarT-associated) NADARs, DarG1-like antitoxin NADARs are still their closest relatives (Fig 4A), suggesting that DarG1 may be their evolutionary origin.

**Figure 4:**
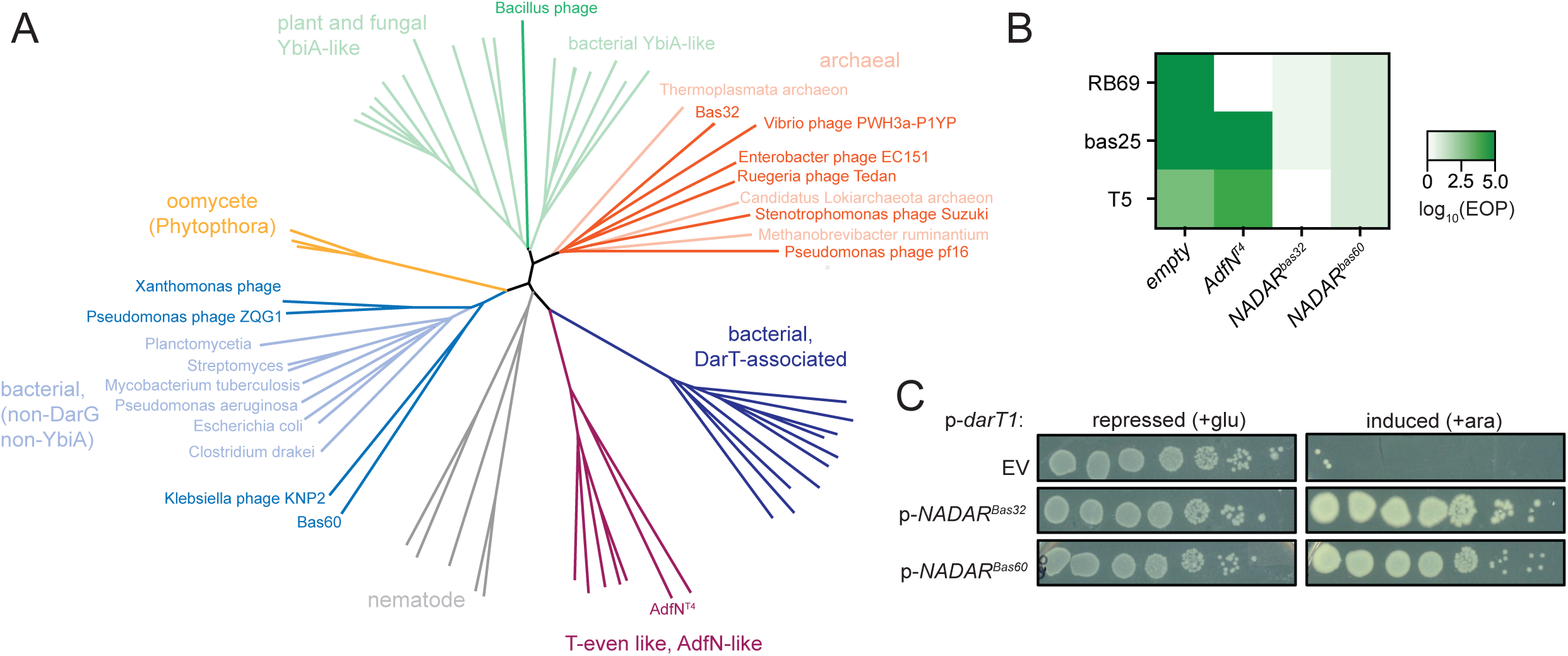
Diverse phage NADARs can counter DarTG1. (A) Representation of the evolutionary relationships among NADAR superfamily members, with phage NADARs indicated. Labels refer to the NADAR type or group of organisms in which a particular NADAR homolog is encoded. See supplemental file for full protein alignments and accession numbers. (B) EOP of indicated phages between DarTG1 and EV hosts, either control (EV) or expressing the indicated NADAR proteins. AdfN data from Figure 3F is replotted for comparison. (C) Cfus of *E. coli* expressing *darT1* and the indicated NADAR protein under *darT1* repressing (+0.2% glucose) or inducing (0.2% arabinose) conditions. Data are representative of at least 3 independent experiments.

We were surprised that no phage NADARs had been described outside of the T4-like phages. To gain insight into whether NADAR proteins have been co-opted on multiple occasions and might be more widespread among phages, we performed a more extensive search for phage NADARs by seeding PSI-BLAST searches with NADARs from other branches of the tree and limiting the search to viruses. These efforts revealed numerous NADAR homologs outside of the T4-like viruses, including in very distantly related phages (Figure 4A). These results prompted us to perform searches of genomes from our phage collection to see if we might have other NADAR-encoding phages on hand. Our initial blastp and PSI-BLAST searches with AdfN did not reveal any homologs; however, a domain enhanced lookup time accelerated search (DELTA-BLAST), which searches against a set of conserved domains, did identify two distinct NADAR family proteins in four phages outside of the *Tevenvirinae* within the BASEL phage collection^23,35^. The Bas32 and Bas33 phages, *Markadamsvirinae* closely related to T5 phage, both encode identical NADAR proteins (hereafter referred to as NADAR^Bas32^), and Bas60, Bas61 and Bas62, three *Vequintavirinae,* encode a second distinct NADAR protein (hereafter NADAR^Bas60^). A phylogenetic analysis of these predicted NADAR proteins revealed that, as expected, they do not cluster with AdfN^T4^ and the other previously identified phage NADARs (Figure 4A)^17^. In a phylogenetic analysis including the NADAR proteins identified in this previous study^17^, in addition to a subset of representative phage NADARs we identified, we found that NADAR^Bas32^ more closely resembles two archaeal NADARs, while NADAR^Bas60^ more closely resembles a group of non-DarT associated, non-YbiA-like bacterial NADARs of unknown function (Figure 4A). These data suggest that phages have co-opted NADARs on multiple occasions from different sources.

We next asked whether the NADAR^Bas32^ and NADAR^Bas60^ are also enzymatically active and able to reverse DarTG1 defense. We hypothesized that because the phages that encode these NADARs are not known to have modified DNA, these NADARs might display broader, less specific activity than AdfN and its relatives. Indeed, ectopic expression of both NADAR^Bas32^ and NADAR^Bas60^ increased phage replication of 3 DarTG1-sensitive phages in the presence of DarTG1 by several logs (Figure 4B), but did not restore replication of a phage blocked by DarTG2 (Figure S3B). Further, both of these NADARs could neutralize the DarT1 toxin in the absence of phage infection (Figure 4C). These results demonstrate that all phage NADARs tested have guanine-specific DNA ADP-ribosylglycohydrolase activity, and, in contrast to AdfN ^T4^, the NADAR^Bas32^ and NADAR^Bas60^ proteins, which are from other phage groups and were likely independently co-opted by phages, are not specific to the DNA from their host.

## Discussion

In this study, we describe the discovery of an enzymatic phage counter-defense element that reverses the activity of the DarT1 toxin and neutralizes DarTG1-mediated phage defense. This protein, AdfN, has a fold similar to the native DarG1 antitoxin and functions in a similar manner via its DNA ADP-ribosylglycohydrolase activity, thus representing to our knowledge the first example of a lytic phage encoding an orphan antitoxin for counter-defense. This activity enables phages to replicate in the presence of DarTG1 by reversing DarT1-mediated DNA ADP-ribosylation and thereby detoxifying their DNA. A single nucleotide insertion in RB69, leading to an early stop codon, resulted in a non-functional AdfN and leaves RB69 uniquely susceptible to DarTG1 defense. Unlike another recently described phage NADAR protein^36^, these proteins require enzymatic activity and thus represent one of only a few examples of enzymatic phage counter-defenses^2^. A unique feature of such a strategy is that a NADAR element can remove ADP-ribose from guanine on DNA regardless of the precise structure of the toxin that placed it there, potentially making for a more versatile type of counter-defense. The phylogenetic relationship between AdfN and DarG1 suggest that phage have potentially co-opted this NADAR protein from the bacterial defense system. Intriguingly, we find NADARs carried by additional, unrelated phages; while these proteins are all able to reverse DarTG1, they belong to other clades of the NADAR superfamily and were likely horizontally acquired in distinct evolutionary events. By revealing the biological function of another subset of the NADAR superfamily, this discovery enables the study of these proteins in their native context during phage infection and underscores the importance of ADP-ribosylation in predator-prey interactions.

While AdfN is related to DarG1, it is also distinct from the native antitoxin in two aspects. First, AdfN and most of the other phage NADARs lack the N-terminal extension that is characteristic of DarT-associated NADARs like DarG1, and whose function is not known. Even when alignments are generated lacking this N-terminal extension, the DarT1-associated NADARs still form a distinct cluster, though the AdfN NADAR cluster is most closely related to these DarG NADAR proteins (Figure 4A). A second distinction between these proteins is that AdfN^T4^ functions most efficiently in the context of *Tevenvirinae* infection: we saw no rescue of DarT1 toxicity in uninfected *E. coli* (Figure 3E), and ectopic AdfN production provides only partial rescue to phages outside of the *Tevenvirinae* from DarTG1 defense (Figure 3F). There are two potential reasons for this specificity: either the enzyme prefers the modified, hydroxymethylated DNA that is found in this group of phages, or these phages encode another factor that is required for full activity. We favor the former model as NADARs generally seem to function as single domain proteins, and because AdfN is able to remove ADP-ribose from unmodified DNA *in vitro* (Figure 3D)^30^. We speculate that the hydroxymethylated DNA stabilizes the interaction between AdfN and its DNA substrate, but the structural basis of this specificity will be an interesting topic of future studies. The AdfN-like NADARs have all replaced an otherwise conserved glutamic acid residue with a histidine; this substitution may have altered specificity of this protein. Together with the fact that DarT1-associated NADARs are most closely related to the AdfN family of NADARs, our results suggest that phages may have co-opted a bacterial antitoxin and then domesticated it, perhaps as a strategy for preventing its horizontal acquisition by competing phages that do not share the same DNA modifications. It also underscores that while DarT1 can act on both phage and host DNA, the primary target of DarT1 in the context of a phage infection is the phage DNA.

Remarkably, around 100 genes in the model T4 phage, which has been studied for many decades, still have no known function. We have now found that a second of these previously hypothetical T4 genes is involved in anti-DarTG1 counter-defense. Further, AdfN homologs are found across the *Tevenvirinae* superfamily, not just in *E. coli* T-even-like phages. Why do the *Tevenvirinae* encode multiple anti-DarT1 defenses? We found that surprisingly, AdfA does not play a major role in DarT1 neutralization for T4, as deletion of *adfA* from T4 has no phenotype, even in phages lacking *adfN* (Fig S3A). We have not further examined other differences between the evolved, functional AdfA from RB69 (AdfA^RB69^(R164H)) and the AdfA homologs in the rest of this group; we think it likely that some of the other differences in these proteins, outside of position 164 (Figure S1), may reduce the ability of AdfA^T4^ to neutralize this specific DarT1. It thus appears that T-even phages have accumulated layers of anti-DarTG counter-defenses that can help protect against the suite of varying DarTG systems that they might encounter, with the T4 AdfA variant likely neutralizing some other DarT homolog similar to the model proposed for AdfB^22^. Notably, neither AdfA nor AdfN – both anti-DarTG1 counter-defenses – counter the DarTG2 defense system, as a T4 phage with both AdfA and AdfN deleted is still resistant to DarTG2 defense (Figure S3A). Yet these phages are all DarTG2 resistant, suggesting that these phages must also have a strategy for countering DarTG2. That such a high proportion of the severely size constrained genomes of *Tevenvirinae* is devoted to countering DarTG systems indicates that these systems have exerted a strong selective pressure on this group of phages. The apparent adaptation of AdfN for the T-even phages further underscores the long evolutionary history between T-even phages and DarTG systems. It is also possible that a subset of these counter-defenses may also detoxify other types of yet, undiscovered DNA ADP-ribosylating phage defenses.

The non-AdfN-like phage NADARs we discovered primarily cluster in two areas of the phylogenetic tree: a subset with archaeal NADARs, and a second, larger set with a group of non-YbiA, non-DarG1-like bacterial NADARs. We see only a single phage NADAR with distant relatedness to the large YbiA-like group that is found in both plants and bacteria. YbiA-like NADARs in both bacteria and plants appear to be housekeeping proteins that have been shown to play a role in riboflavin metabolism^37^. We tested a representative of both the archaeal-type and bacterial-type NADAR, and found both to exhibit DNA ADP-ribosylglycohydrolase activity, further validating that the NADAR superfamily, outside of YbiA-like proteins, share similar activity, despite their low sequence homology.

Orphan antitoxins have rarely been described in bacterial genomes, though this may be due to bioinformatic challenges in their identification^38^. The DarG antitoxin is a type IV antitoxin with enzymatic activity and a conserved fold^18^, making it relatively easy to identify bioinformatically. In contrast, type II TA systems, which have been studied in most details, consist of antitoxins that neutralize their cognate toxins via a direct interaction and occlusion of the active site. These antitoxins are typically unstructured when unbound to their cognate toxin, and thus lack a distinctive fold or structural homology^39–42^. They are also highly specific to their cognate toxin, with little ability to cross-neutralize even closely related toxins^43^. However, the idea of chromosomal antitoxins – even those associated with a functional intact TA system – functioning as antitoxins to an incoming TA system has been proposed. In this model, the chromosomal antitoxin would function as an anti-addiction module to neutralize the toxin of a plasmid encoded TA system, thereby enabling the loss of the plasmid, though there is limited experimental evidence of such interactions^26,44^.

However, an analogous example of an orphan antitoxin, or immunity protein, functioning to neutralize a toxin, can be found in the field of interbacterial antagonism, and could provide a clue regarding the function of the stand-alone bacterial NADARs. Interbacterial toxins delivered by a type VI secretion system are encoded adjacent to a cognate immunity protein that protects bacteria from self-intoxication. Such immunity proteins have been found in the absence of a cognate toxin and these have been experimentally demonstrated to provide defense from bacterial antagonism^45^. One intriguing possibility is that the abundant stand-alone NADARs found in bacteria may play a role in protecting their hosts from ADP-ribosylation resulting from a toxin delivered by a bacterial competitor. Indeed, a family of RNA ADP-ribosyltransferases delivered by the type VI secretion system have been described^46^; it seems entirely possible that there are as-yet undiscovered guanine-targeting DNA ADP-ribosyltransferases delivered in a similar manner. Further, two additional guanine targeting DNA ADP-ribosyltranferases have been identified in eukaryotes (pierisins in butterfly larvae, and the CARP-1 toxins in clams)^47^. Perhaps some bacterial NADARs, like their phage counterparts, are maintained in bacterial genomes to neutralize a eukaryotic defense mechanism. It is exciting to speculate that potentially counter-defenses, similar to what has been found for bacterial immune mechanisms, may also be conserved across domains of life^48–50^.

## Acknowledgements

This work was supported by an NIAID DP2 (DP2AI177955). We thank members of the LeRoux lab for helpful discussions and comments on this manuscript. We thank Sriram Srikant in the Laub lab for the T4 Δ*agt* Δ*bgt* strain and for helpful discussions.

## Author contributions

Conceptualization, **M.L**.; Methodology, **M.L.;** Investigation, **M.L**.**, A.J., N.C.**; Writing**, M.L.**; **Visualization, M.L.;** Funding Acquisition, **M.L**.; Formal Analysis, M.L.; Supervision, M.L.

## Methods

### Strains and growth conditions

All bacterial and phage strains are listed in Table S1. *Escherichia coli* was grown at 37 °C in LB medium for routine maintenance and cloning. Phages were propagated by infecting *E. coli* MG1655 cultures of OD ∼0.1-0.3 at an MOI of 0.1 and incubating with aeration at 37 °C. Following clearing, any remaining cells were pelleted by centrifugation and lysates were filtered through a 0.22 µM filter. Media for selection or plasmid maintenance were supplemented with carbenicillin (100 µg/mL), chloramphenicol (20 µg/mL), or kanamycin (30 µg/ml) as necessary unless otherwise indicated. Induction of ectopic expression were effected with anhydrous tetracycline (aTC) (10 ng/µL) or arabinose (0.2% w/v). Media were supplemented with glucose (0.4%) to repress DarT1 toxin production as needed.

### Bioinformatics

For AdfA, a blastp search was run on the genomes of *Tevenvirinae* phages found within the Basel phage collection as well as T2, T4, T6, and RB69. Proteins that were >75% of full length were aligned using Muscle algorithm in Geneious. PSI-BLAST searches were performed with NADAR proteins from each major cluster of the phylogenetic tree as described in a previous study^17^ with results limited to “Viruses” and representative phage NADAR amino acid sequences were selected. To identify NADARs in the BASEL collection, a DELTA-BLAST search was performed in NCBI selecting the BASEL phages in the “organism” field. In addition to these newly identified phage NADARs, the entire set of proteins described in Schuller et al. (2023) were obtained and the NADAR domains were trimmed in each case. The resulting amino acid sequences were aligned using the Muscle algorithm in Geneious. The phylogenetic tree was created in SplitsTree with the Neighbor-Joining algorithm. A full protein alignment is provided as supplemental data.

### Phage deletions

Deletions were made in phages using a recombination template and counter-selection with CRISPR-Cas13 as described previously^25^. Recombination templates consisted of ∼250 bp of flanking regions with a small scar region were cloned via Gibson assembly into either pBA701 or pSSRescue, a plasmid that contains strong terminators flanking the homology regions to facilitate cloning by reducing the expression and thus toxicity of phage DNA in bacterial cells (kind gift of Sriram Srikant, Laub lab). The phage being modified was propagated on the recombination template and the resulting lysate was then propagated on a strain encoding the CRISPR-Cas13 plasmid with a guide targeting the region being deleted. Guides were initially tested for restriction and either used with leaky expression (if induction was toxic) or induced at 10 ng/µL aTC. A dilution of the resulting lysate was spotted on a top agar plate made with the CRISPR-Cas13 bearing strain, and individual plaques were checked by PCR for the deletion. All phage deletion strains were confirmed by PCR and sequencing.

### Phage crosses

Phages being crossed were mixed at a 1:1 ratio and MG1655 cells were infected at an MOI of 0.1. Upon clearance of the culture, the lysate was centrifuged and filtered through a 0.2 µM filter to remove any remaining cells. The titer of the resulting lysate was determined, and then an overnight culture of *E. coli* containing the CRISPR-*cas13^dmd^* and pBR322-*darTG1* vectors were infected at an MOI of 0.1. For the guide used in these experiments targeting the *dmd* gene, the CRISPR-Cas13 targeting of T2 and T4 was found to be nearly complete with no added aTC; further inducing this construct led to toxicity of bacteria. Cleared cultures were centrifuged, the supernatant filtered, and 10-fold serial dilutions of the resulting lysate were spotted on a top agar plate made with the same selection strain. Individual plaques were picked and propagated on the selection strain. DNA from phage was extracted from these propagations with the Norgen Biotek Phage DNA Isolation kit (Norgen Biotek) and genomic phage DNA was sequenced via short-read Illumina sequencing by SeqCoast. Sequencing data were deposited to SRA (PRJNA1120483).

### Plasmid construction

CRISPR-Cas13 guide plasmids were constructed via Golden Gate assembly as described previously^25^. Briefly, complementary oligos targeting a 31 base region were ordered to include BsaI sites. Oligos were annealed by heating to 98°C and slowly cooling, then treated with PNK to phosphorylate ends, and ligated to pBA559 digested with BsaI-HF. The recombination template for the deletion of *adfA* from RB69 was created using Golden Gate assembly using a gene fragment (Twist Biosciences) with ends that include a BbsI cut-site and ligated into pBA707 as described^25^. All other plasmids were created using Gibson assembly with NEB HiFi DNA Assembly MasterMix (NEB). Vector backbones and inserts were generated via PCR using primers listed in Table 2. Insertions were verified by Sanger or Nanopore sequencing.

**Table 1:**
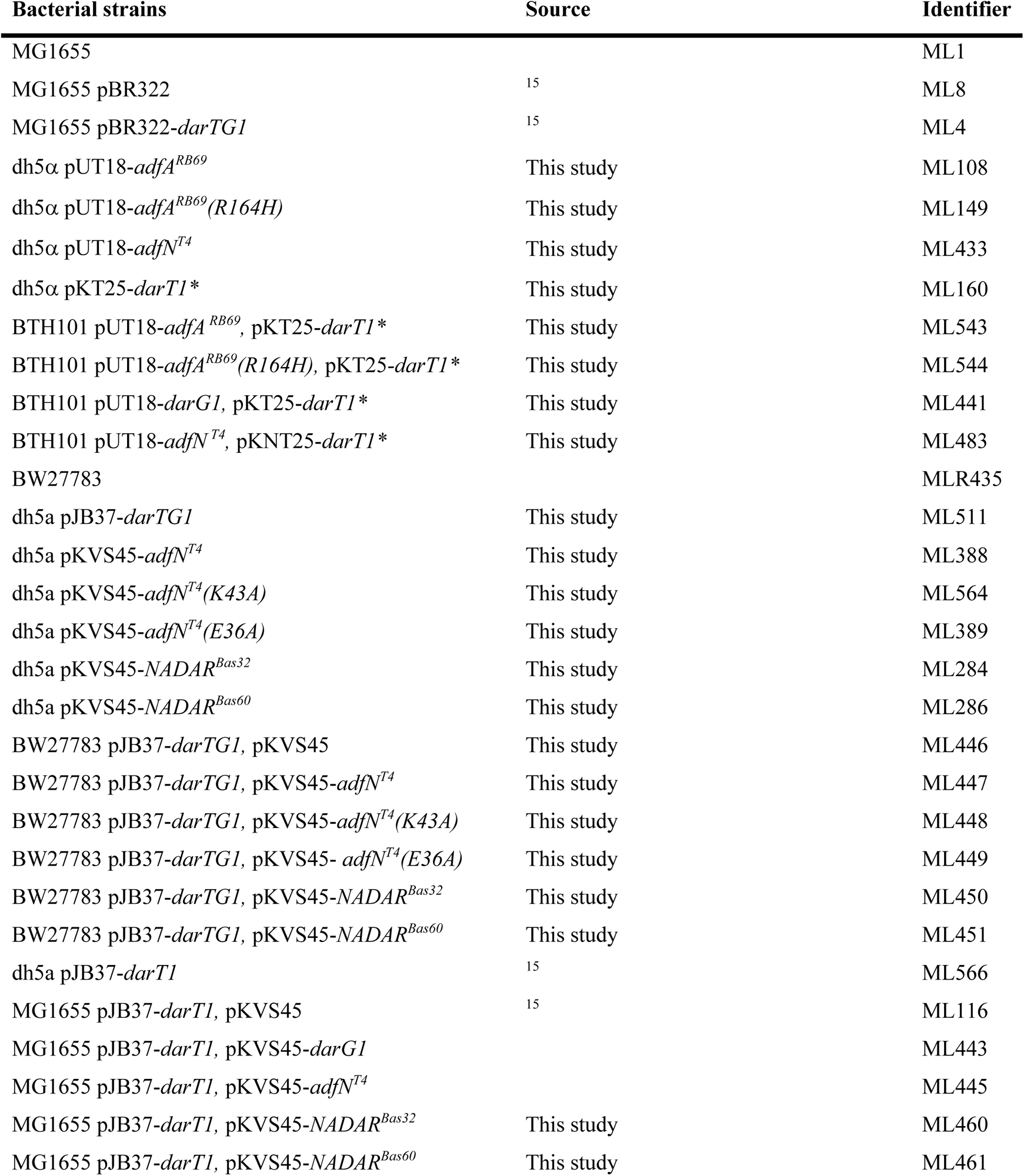
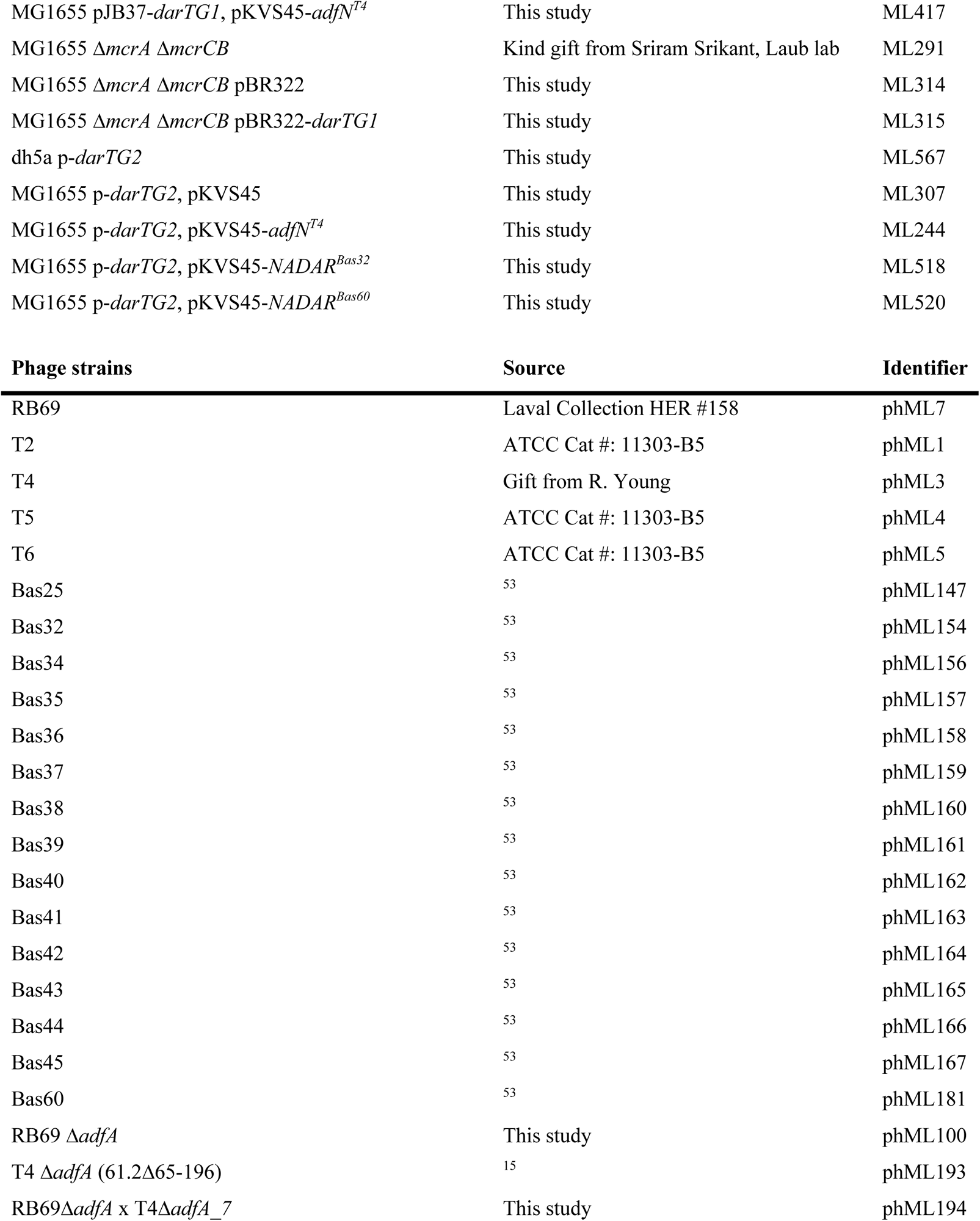
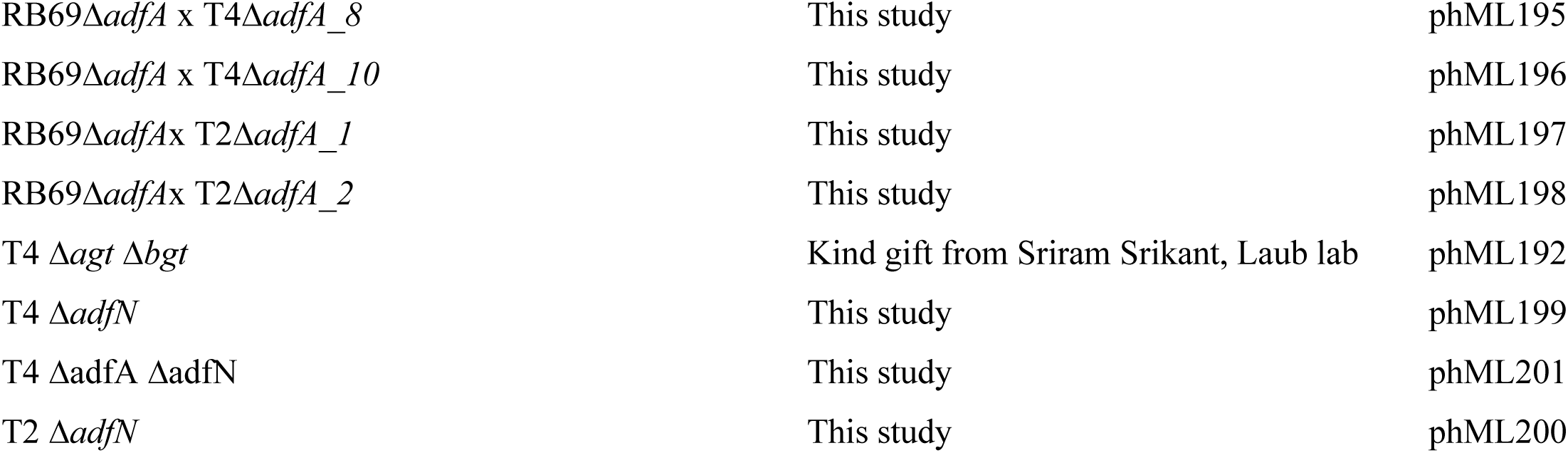
Strains used in this study.

**Table 2:**
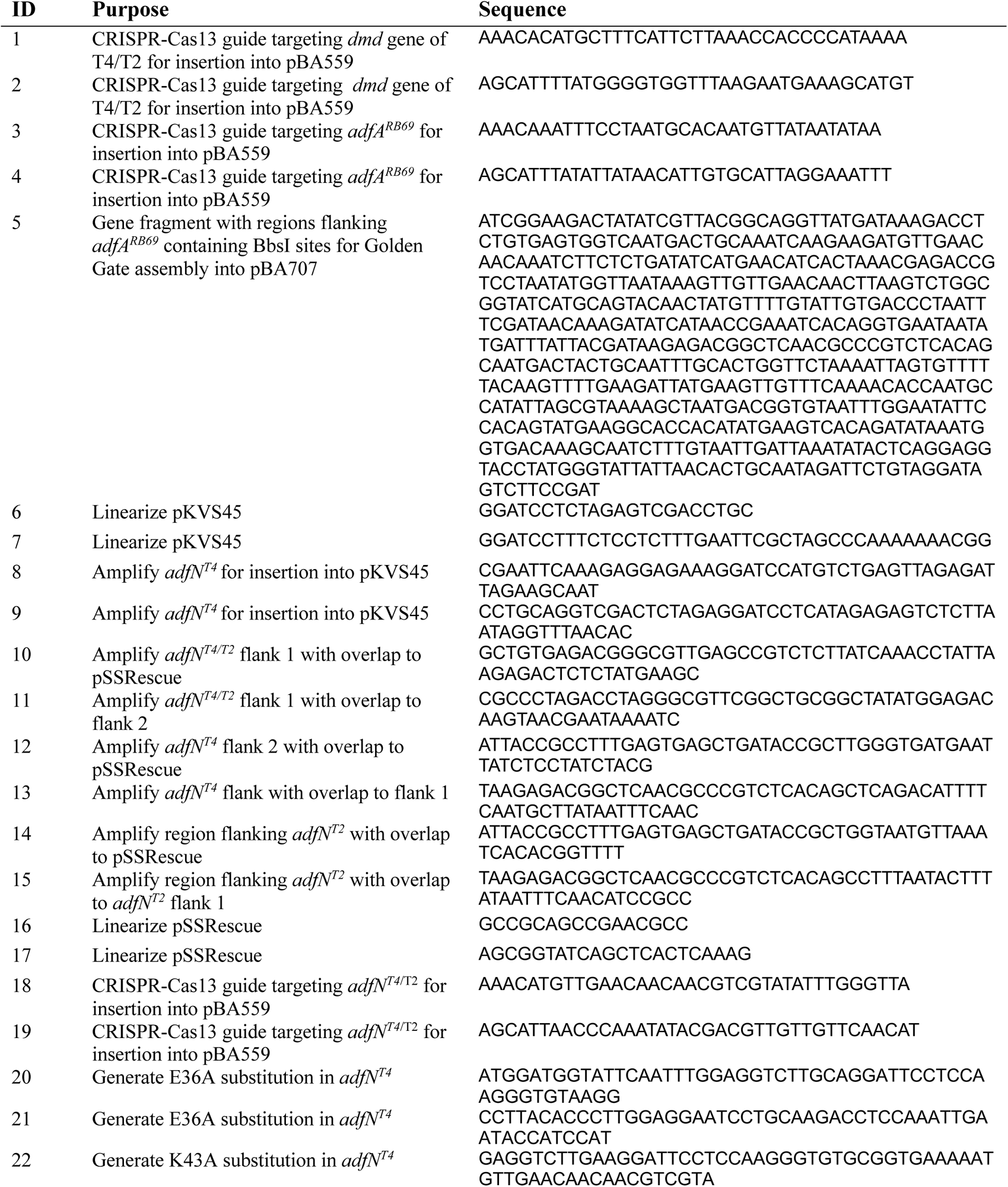
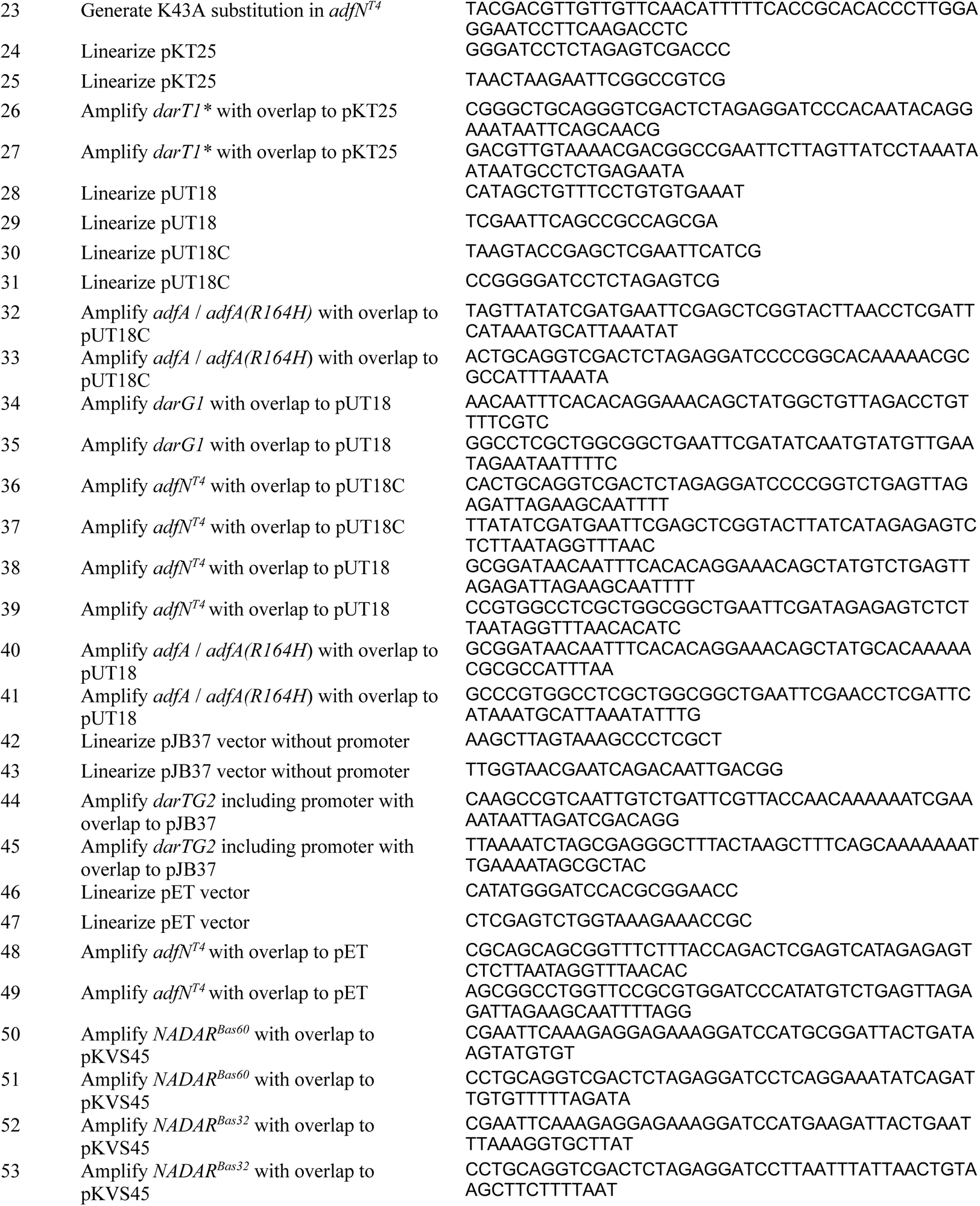
Primers used in this study.

### Plaque Assays

Plaque assays were performed using LB medium plates with 1.2% agar plates. Top agar was prepared with melted 0.5% agar-LB and combined with overnight cultures 1:200 in a 4 mL volume. Both the plates and the overlay were supplemented as needed for different experiments as described below. Two microliters of ten-fold serial dilutions of phages were spotted onto the overlaid plates and incubated at 37 °C. All plates were imaged with an Epson Perfection V600 Photo Scanner following overnight growth. For Figure 1A, DarTG1 defense against the full *Tevenvirinae* panel was assessed with the pBR322-*darTG1* vector in which *darTG1* is expressed from its native promoter^15^. The same conditions were used in Figure 2B (testing *adfN* deletions in T2 and T4). For rescue of phage with ectopic counter-defense genes, *darTG1* was expressed from the pJB37 vector (SC101 origin, arabinose promoter) and the putative counter-defense gene was cloned into pKVS45 (P15a origin, tet promoter), and media were supplemented with aTC (10 ng/uL) and L-(+)-arabinose (0.2%). Expression of DarTG2 experiments were performed with *darTG2* expressed from its native promoter on a low-copy plasmid (SC101 origin). Because DarTG2 defense is more potent under conditions that reduce bacterial growth rates^15^, sub-MIC chloramphenicol (2 ug/mL) was added to plates for all DarTG2 plaque assays. All experiments were repeated independently at least three times.

### Bacterial two-hybrid assays

The bacterial adenylate cyclase two-hybrid system was used to assay protein interactions^51,52^. To assess genes of interest, the genes were cloned onto the pKT25, pKNT25, pUT18, and pUT18C vectors and were fused to the 3’ or 5’ ends of the T18 and T25 fragments of the *Bordetella* adenylate cyclase and transformed into *E. coli* BTH101. The transformants, a combination of one T18 plasmid with one T25 plasmid, were grown overnight and spotted on LB agar plates supplemented with X-gal (10 mL/L). Plates were left to develop at 30 °C for 24 hours and imaged on an Epson photo scanner.

### Toxin neutralization assay

Each strain contained the toxin vector, pJB37-*darT1*, and a gene of interest on the other vector, pKVS45. The strains were grown overnight in LB medium with antibiotics and glucose (0.4%), then tenfold serially dilutions were spotted on plates supplemented with spectinomycin, anhydrotetracycline (10 ng/uL, to induce the putative counter-defense element), and either arabinose (0.2%) or glucose (0.4%) to induce or repress *darT1*, respectively. The plates were incubated at 37 °C and imaged after 16 hrs on a Epson V600 photo scanner.

### Protein purification

AdfN and AdfN(K43A) were expressed from pET vectors with a 6x-His tag in Rosetta 2 (DE3) cells (Millipore Sigma). For expression, 500 mL cultures were grown at 37°C until an OD_600_ of 0.5; cultures were induced with 0.5 mM IPTG and moved to 30 °C overnight. Cultures were centrifuged and pellets frozen at -80°C until lysis. Cells were resuspended in lysis buffer (50 mM HEPES, 500 mM NaCl, 5% v/v glycerol, 20 mM imidazole, 10 µM DTT, 10 units benzonase, EDTA-free protease inhibitor) and lysed via two passes through a Constant Systems Cell Disruptor at 30 psi. Resulting lysates were clarified by centrifugation at 10,000 x g for 2 hours. Clarified lysates were loaded to an equilibrated column of Nickel-NTA resin (Goldbio), washed with 10-15 column volumes of wash buffer (50 mM HEPES, 500 mM NaCl, 5% v/v glycerol, 10 µM DTT), and eluted with a stepwise increase in imidazole (100-400 mM). Fractions were analyzed by SDS-PAGE and Coomassie, concentrated in a 10 kD protein concentrator column (Pierce), and transferred to storage buffer. Purified protein was snap frozen in liquid nitrogen and stored at -80°C.

### ADP-ribose glycohydrolase assays

Overnight cultures of *E. coli* dh5a pJB37-*darT1* were grown with 0.4% glucose, then diluted 1:100 with fresh, glucose-containing media until an OD_600_ of ∼0.3. A serial dilution of the culture was also spotted onto an arabinose containing plate to confirm that escape mutants do not make up a large proportion of the culture, which arise quickly in these cultures. Upon reaching the desired density, the cultures were washed and released into media containing 0.2% arabinose. After an additional 45 min of growth, the culture was collected, centrifuged, and cell pellet was stored until the arabinose-spotted culture can be verified for a lack of growth. DNA was extracted from the pellet using the Qiagen PureGene DNA extraction kit, and sheared with a BioRuptor sonicator for 5 x 30 seconds at maximum intensity to improve DNA solubility. DNA concentration was assessed using a Nanodrop. One µg of DNA was then incubated in a reaction containing 10 µM AdfN or AdfN^E36A^ in a buffer consisting of 10 mM HEPES, 5% glycerol, and 5 mM NaCl. The DNA was purified with a Zymo DNA Clean and Concentrator kit and spotted on a nitrocellulose membrane. The resulting membrane was cross-linked in a Spectrolink XL-1000 UV Crosslinker with the Optimal Crosslink setting (1.2 x 10^5^ µJ/cm^2^), blocked in 5% milk tris-buffered saline supplemented with 0.5% v/v Tween-20 (TBS-T), and incubated with a poly/mono ADP-ribose antibody (D9P7Z, Cell Signaling) diluted 1:1000 in 5% milk TBS-T for 2 hrs at room temperature or overnight at 4 °C. The membrane was washed 3 x 5 min in TBS-T, then incubated with a goat anti-rabbit-HRP conjugated antibody (ThermoFisher) at 1:25000 in 5% milk, before being washed and developed with SuperSignal West Femto Reagent (ThermoFisher) and imaged on a BioRad ChemiDoc system using the chemiluminescence setting. Following imaging, the membrane was briefly washed in water, then incubated in a solution of 0.1% (w/v) methylene blue in 0.5 M sodium acetate, pH 5.2 for 2 min, washed with distilled water to destain for 2 min, then dried and imaged.

---

**Figure S1:**
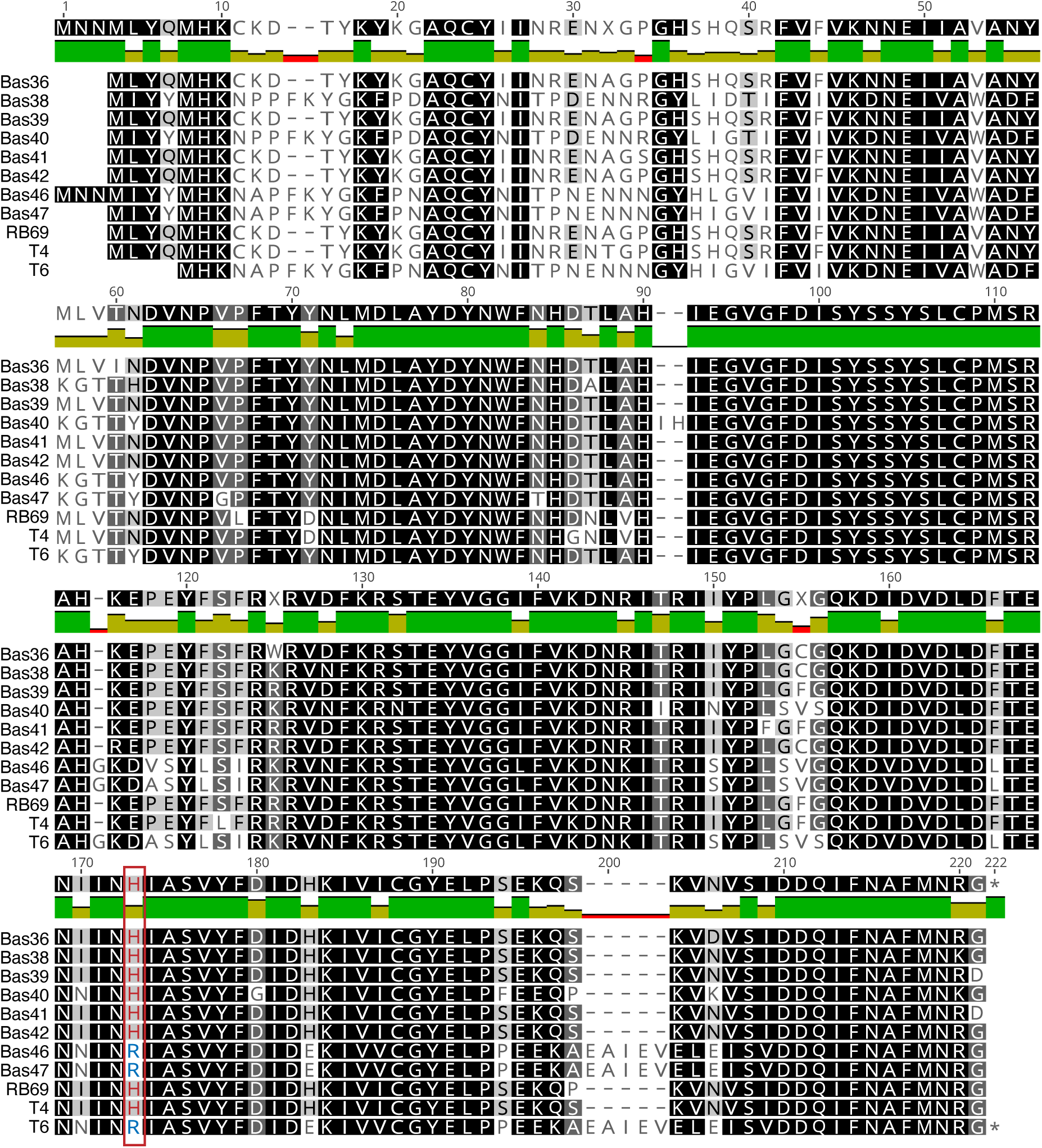
AdfA varies among *Tevenvirinae*. Amino acid alignments of AdfA homologs identified in the set of *Tevenvirinae.* The position shown to correlate with DarT1 neutralization is highlighted with a red box.

**Figure S2:**
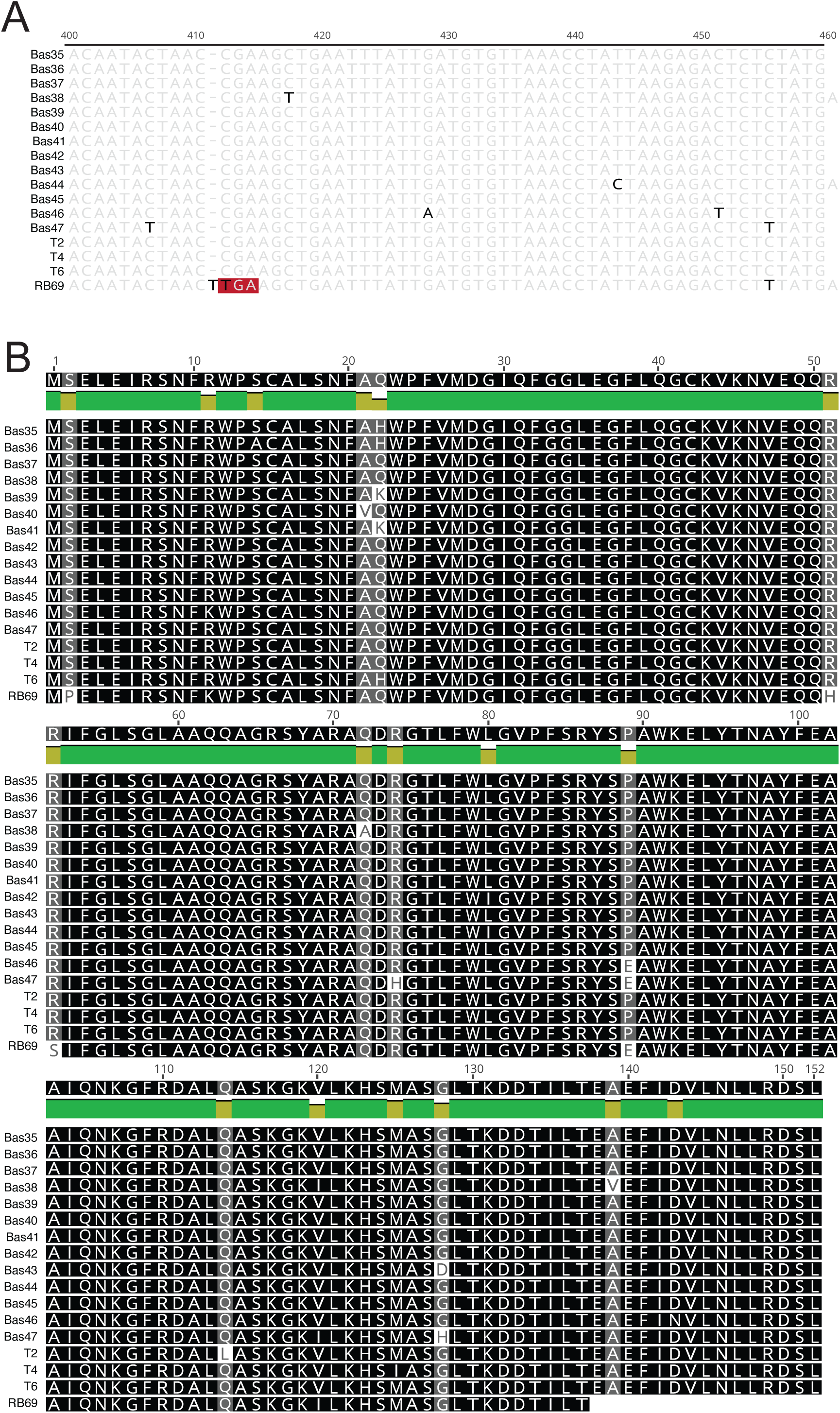
AdfN is highly conserved among the *Tevenvirinae* phages but mutated in RB69. (A) Nucleotide alignment of the last 60 bases of *30.3* from the *Tevenvirinae*. Mutations are highlighted in black, and the early stop codon in RB69 is highlighted with a red box. (B) Amino acid alignment of AdfN (gp30.3) homologs from the set of *Tevenvirinae*.

**Figure S3:**
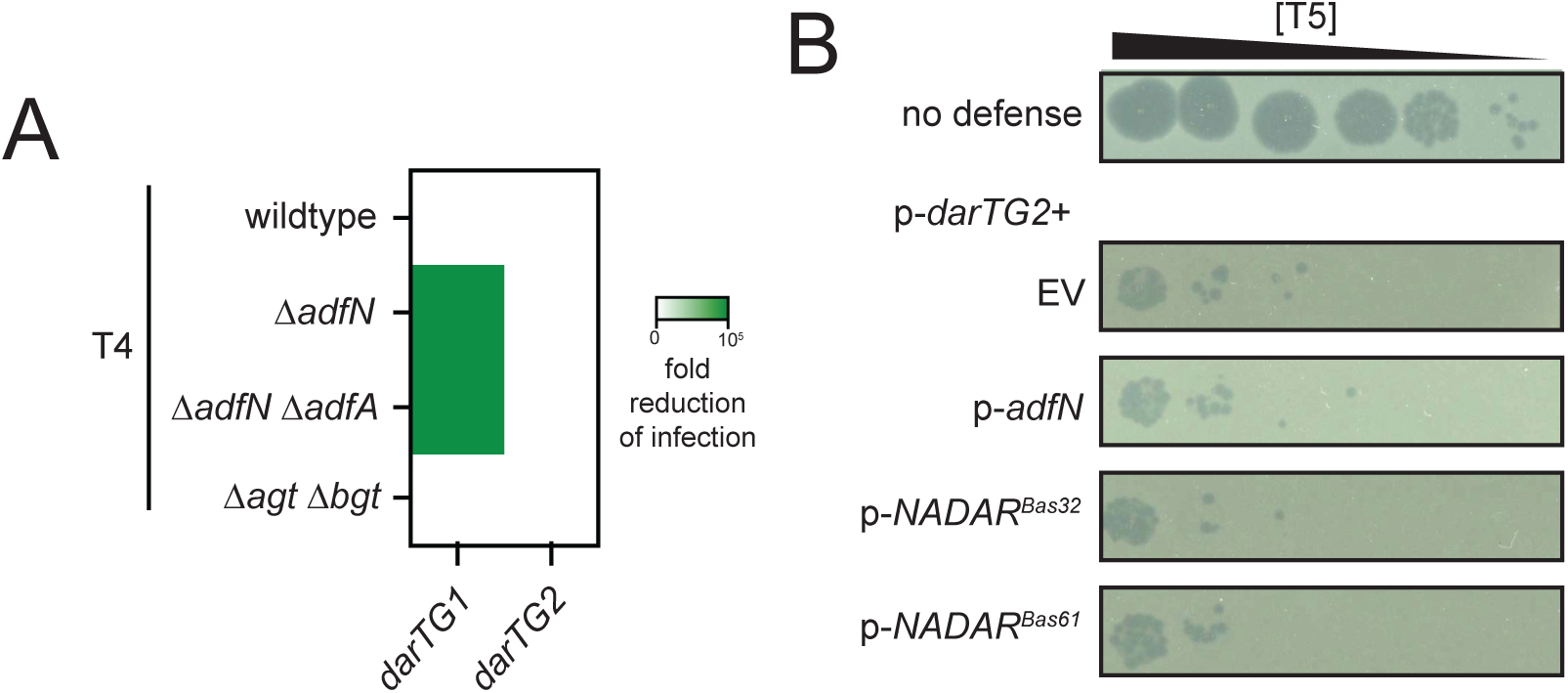
Neither AdfA nor AdfN counter DarTG2. (A) EOP of the indicated T4 strain on *E. coli* expressing *darTG1* or *darTG2*, compared to an empty vector. (B) EOP of T5 on *E. coli* expressing *darTG2* from its native promoter in addition to the indicated NADAR domain protein expressed ectopically from the tet promoter.

